# Drug-tolerant idling melanoma cells exhibit theory-predicted metabolic low-low phenotype

**DOI:** 10.1101/809889

**Authors:** Dongya Jia, B. Bishal Paudel, Corey E. Hayford, Keisha N. Hardeman, Herbert Levine, José N. Onuchic, Vito Quaranta

**Author notes:** These authors contributed equally (Dongya Jia, B. Bishal Paudel). These authors contributed equally (Herbert Levine, José N. Onuchic, Vito Quaranta). Correspondence to: Vito Quaranta, Herbert Levine, Jose Onuchic. **Author Contributions:** Conceptualization: D.J., B.B.P.; Data Acquisition, Analysis, and Interpretation: D.J., B.B.P, K.N.H., C.E.H., V.Q., H.L., & J.O. Writing: D.J., B.B.P., V.Q., H.L., J.O. All authors reviewed and revised the manuscript.

## Abstract

Cancer cells adjust their metabolic profiles to evade treatment. Metabolic adaptation is complex and hence better understood by an integrated theoretical-experimental approach. Using a minimal kinetic model, we predicted a previously undescribed Low/Low (L/L) phenotype, characterized by low oxidative phosphorylation (OXPHOS) and low glycolysis. Here, we report that L/L metabolism is observed in *BRAF*-mutated melanoma cells that enter a drug-tolerant “idling state” upon long-term MAPK inhibition (MAPKi). Consistently, using publicly available RNA-sequencing data of both cell lines and patient samples, we show that melanoma cells decrease their glycolysis and/or OXPHOS activity upon MAPKi and converge toward the L/L phenotype. L/L metabolism is unfavorable for tumor growth, yet supports successful cell division at ~50% rate. Thus, L/L drug-tolerant idling cells are a reservoir for accumulating mutations responsible for relapse, and it should be considered as a target subpopulation for improving MAPKi outcomes in melanoma treatment.

## Introduction

Cancer cells can adapt their metabolic profiles to survive drug treatment. Traditionally, the Warburg effect or aerobic glycolysis was regarded as the dominant metabolic phenotype in cancer [1]. Recently, mitochondrial OXPHOS has been shown to also play a critical role, especially during metastasis and drug-resistance [2]. Accumulating evidence suggests that cancer cells, in response to external perturbations, can adapt their metabolic profiles by utilizing glycolysis, OXPHOS, or both, referred to as metabolic plasticity [3]. For instance, melanoma cells upon BRAF inhibition (BRAFi) can exhibit enhanced OXPHOS [4,5]. We and others [6,7] have shown that BRAFi is most effective when melanoma cells rely primarily on glycolysis (or forced to utilize glycolysis by depleting mitochondria). Drug-tolerant cancer cells have been shown to have increased dependency on certain substrates such as glutamine [5] and polyunsaturated lipids [8]. Taken together, these results underscore a critical role of metabolic adaptation in cancer cell survival.

To elucidate metabolic plasticity in cancer, we established a modeling framework that couples gene regulation with metabolic pathways (**electronic supplementary material, figure S1**) [3]. Our modeling analysis demonstrates a direct association of the master gene regulators of cancer metabolism AMP-activated protein kinase (AMPK) and hypoxia-inducible factor (HIF)-1 with three major metabolic pathways - glycolysis, glucose oxidation and fatty acid oxidation (FAO). Specifically, the model predicts the existence of a hybrid metabolic state where cancer cells can use both glycolysis and OXPHOS, and a metabolically inactive state where cancer cells exhibit low activity of both glycolysis and OXPHOS, referred to as the “low/low” (L/L) state. The importance of this new state has to date remained uncertain.

It is well-established that bacterial populations can harbor a subpopulation of persister states that exhibit tolerance to antibiotics [9]. Interestingly, persisters seem to have altered metabolism as compared to the drug sensitive cells. Similar to bacterial persisters, several possible forms of drug ‘persisters’ have been described in cancer, including quiescent [10], senescent [11], dormant [12] and drug-tolerant cells [13]. For example, recent studies have implicated the emergence of slow-cycling phenotype as a mode of drug resistance in melanoma [14,15]. Slow-cycling melanoma cells are shown to elevate their OXPHOS to continue to grow slowly, by lengthening their G1 state. Other studies show that the expression of MITF or AXL can define distinct phenotypic states including the drug-resistant state in melanomas [16,17]. These studies suggest that melanoma cells can reside in distinct drug-sensitive states, and the transition among them likely describes how melanoma cells respond to therapy. Interestingly, phenotypic transition, as an escape route, has further been documented in studies, which show a therapy-induced differentiation towards neural-crest phenotype [18]. Recently, we reported that *BRAF*-mutated melanoma cells converge toward a non-quiescent “idling population state” upon longterm MAPKi treatment [19]. In a sense, the idling state defines a form of drug-tolerance since the total cell population does not expand; rather, it is maintained at a constant size (i.e. it idles) by undergoing cell division and death at approximately equal rates within clonal lineages of singlecell derived subclones. Notably, this is different from the slow-cycling phenotype as the idling cells continue to divide at moderate doubling times. Nonetheless, each round of successful cell division brings with it the possibility of accumulating mutations, a gateway to acquired drug resistance under drug selective pressure, and concomitant relapse.

Inspired by the theoretical predictions and the analogy to bacterial persisters, we investigated the possibility that the L/L phenotype could characterize the drug-tolerant idling state. Here, we report that the *BRAF*-mutated melanoma, SKMEL5 cells can indeed repress both their glycolysis and OXPHOS activity when they transition into the idling state as a response to therapy. Furthermore, using the RNA-sequencing (RNA-Seq) data of human melanoma M397 and M229 cell lines and melanoma patient samples from Gene Expression Omnibus (GEO), we show that melanoma cells consistently decrease their glycolysis and/or OXPHOS activity and exhibit a convergence toward the L/L phenotype upon long-term MAPKi treatment, as characterized by the AMPK/HIF-1 signatures [20] and the metabolic pathway scores [3]. The residual melanoma tumors may be composed of a significant fraction of drug-tolerant idling cells. We further use our model to explore conditions that promote the L/L phenotype and show that high HIF-1 degradation or low mtROS production can significantly enhance the L/L phenotype. In summary, the L/L metabolic phenotype, predicted by modeling and validated by experiments, is clinically relevant and can be a potential bottleneck for melanoma treatment.

## Materials and Methods

### 1. The mathematical model of metabolism

We utilize our previously developed mathematical model of metabolism [3,20] to analyze the emergence of the L/L phenotype. In brief, the generic form of the differential equation representing the temporal dynamics of x (pAMPK, HIF-1, mtROS or noxROS) is

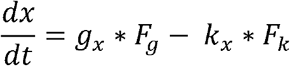

where *g_x_* and *k_x_* represent the basal production and degradation rates of *x* respectively. *F_g_* and *F_k_* represent the regulation of the production and degradation of *x* by others respectively. Considering the chemical reaction happens much faster relative to gene regulation, we assume the metabolic pathways reach an equilibrium state at certain levels of pAMPK and HIF-1. The details of all equations can be found in the **electronic supplementary material, §S1**. The values of all parameters can be found in **electronic supplementary material, tables SS2-3**.

### 2. The AMPK/HIF-1 signature

Since the active form of AMPK is pAMPK, the mRNA expression of AMPK is not indicative of AMPK activity (*23*). HIF-1 is often stabilized in cancer due to the commonly hypoxic condition faced by cancer cells. The mRNA expression of HIF-1 is also not indicative of HIF-1 activity. Phosphorylated AMPK (pAMPK) can up-regulate three major transcription factors - PGC-1α, CREB and FOXO, whose downstream genes are used to evaluate the activity of AMPK. HIF-1 is a transcription factor and its downstream genes are used to evaluate its activity. We quantify the AMPK and HIF-1 activity by evaluating the expression of their downstream targets respectively. We perform PCA of the gene expression data of AMPK and HIF-1 downstream targets respectively, from which we assign the PC1s as the signatures. The AMPK and HIF-1 downstream target genes can be found in **electronic supplementary material, table S4**.

### 3. The metabolic pathway scoring metric

We assume that higher activity of the metabolic process would need higher levels of enzymes which function in that process. Higher metabolic pathway scores indicate higher activity of the corresponding metabolic process. With this assumption, we quantify the activity of TCA, glycolysis and FAO by considering the expression of a total of 10 TCA enzyme genes, 8 glycolysis genes and 14 FAO enzyme genes. The metabolic pathway score is defined as follows, 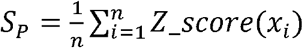 where *x_i_* represents the expression of enzyme gene *i* and *n* represents the total number of genes analyzed for pathway *P* (TCA, FAO or glycolysis). The full list of the enzyme genes can be found in **electronic supplementary material, table S5**. The genes used in the metabolic pathway scoring metric do not overlap with the genes used to construct the AMPK/HIF-1 signatures.

### 4. RNA-Seq of single-cell derived melanoma cells

RNA-Seq of melanoma cell lines (SKMEL5 subclones) was performed as previously reported [21]. Briefly, RNA samples for each subclone, each in triplicate, were collected at three time points (day 0, day 3, and day 8) upon treatment with 8μM PLX4720. All downstream analyses were performed in R (https://www.r-project.org) using the Bioconductor framework (https://www.bioconductor.org). The expression data can be found in **electronic supplementary material, table S6**.

### 5. Seahorse metabolic assays

Seahorse metabolic assays to measure the oxygen consumption rate (OCR) and extracellular acidification rate (ECAR) were performed as previously reported [6]. In brief, SKMEL5 subclones were treated with either DMSO or 8μM PLX4720 for 8 days and plated in 96-well plates (Seahorse Biosciences, Bilerica, MA) at a density of 25,000 cells 24 hours before analysis on the Seahorse XFe 96 extracellular flux analyzer. Mitochondrial oxygen consumption was quantified using the Mito Stress Test kit, and the glycolytic rate was quantified using the Glycolysis Stress Test kit, each according to the manufacturers’ instructions.

## Results

### Modeling-Predicted Metabolically L/L Phenotype in Cancer

We previously developed a metabolic modeling framework that couples the regulatory circuit AMPK:HIF-1:ROS with three major metabolic pathways - glycolysis, glucose oxidation and FAO (**electronic supplementary material, figure S1**). The outputs of the model are the stable steady-state levels of pAMPK, the active form of AMPK, HIF-1 (H), mtROS (R_mt_), noxROS (R_nox_) and the rates of glucose oxidation (G1), glycolysis (G2) and FAO (F). Using the modeling framework, we show that cancer cells can robustly acquire four stable metabolic states “O”, “W”, “W/O” and “L/L” corresponding to an OXPHOS phenotype, a glycolysis phenotype, a hybrid metabolic phenotype, and a metabolically inactive low/low phenotype respectively (**figures 1A-C**). The “O” state is characterized by high AMPK activity and high OXPHOS pathway activity. The “W” state is characterized by both high HIF-1 activity and high glycolysis pathway activity. The “W/O” state is characterized by high AMPK/HIF-1 activity and high OXPHOS/glycolysis pathway activity. The “L/L” state is characterized by low AMPK/HIF-1 activity and low OXPHOS/glycolysis pathway activity (**figure 1C**). The existence of this “L/L” state has until now remained an unverified prediction. Notably, the fractional occurrences of “W” and “O” states within our model are often much larger than those of “W/O” and “L/L” states, indicating the emergence of “W/O” and “L/L” states probably requires specific modifications to the parameters of the baseline cellular network.

**Figure 1.**
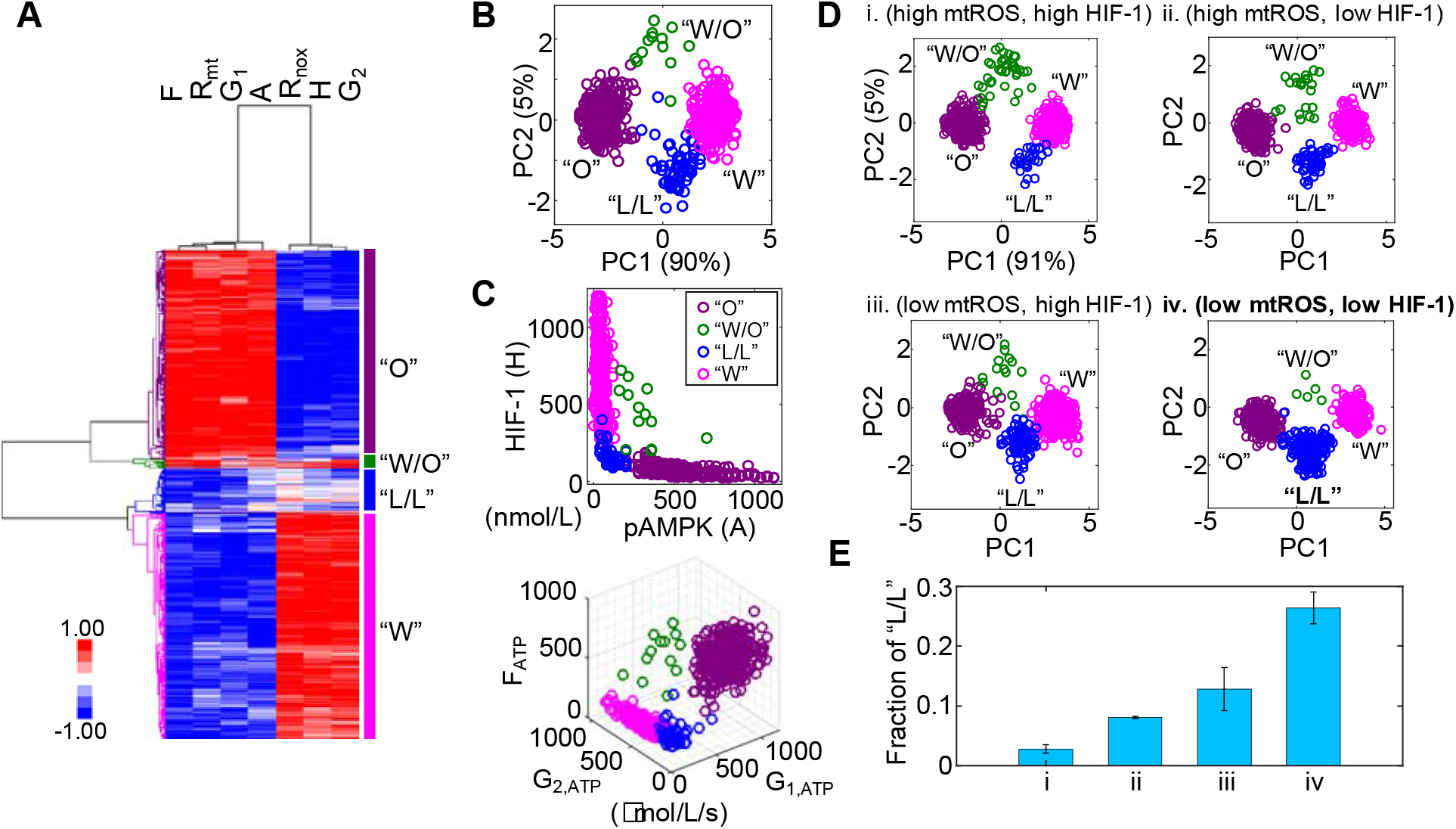
Modeling-predicted metabolically inactive L/L phenotype in cancer. (*A*) Hierarchical clustering analysis (HCA) of the stable-state solutions from 500 sets of parameters. Each row represents one stable-state solution, referred to as one sample here. Each column represents the levels of a regulatory protein, a metabolite or the rate of one metabolic pathway. “F” represents the FAO rate, “R_mt_” represents the level of mtROS, “G_1_” represents the glucose oxidation rate, “A” represents the level of pAMPK, R_nox_ represents the level of noxROS, “H” represents the level of HIF-1 and “G_2_” represents the glycolysis rate. Different colors of the lineages correspond to different clusters. “O” corresponds to an OXPHOS phenotype. “W/O” corresponds to a hybrid metabolic phenotype. “L/L” corresponds to a low/low phenotype and “W” corresponds to a glycolysis phenotype. (*B*) PCA of the clustered samples in (A). Projection of the clustered sample in *A* to the pAMPK and HIF-1 axes (*C, top*), to the G_1, ATP_, G_2, ATP_, and F_ATP_ *axes(C, bottom*). G_1, ATP_, G_2, ATP_, and F_ATP_ represent the ATP production rates of glucose oxidation, glycolysis and FAO respectively. (*D*) Evaluating the effects of HIF-1α degradation and mtROS production on the fractions of different metabolic states. Two values of mtROS production rates 45 mM/min and 30 mM/min are used to simulate “high mtROS” and “low mtROS” respectively. Two values of HIF-1 degradation rates 0.25 h^−1^ and 0.45 h^−1^ are used to simulate “high HIF-1” and “low HIF-1” respectively. 500 sets of parameters were used to collect the stable-state solutions for each scenario. The solutions of each scenario were first normalized using the mean and standard deviation (std) calculated in the scenario (high mtROS, high HIF-1), clustered and projected onto the PC1 and PC2 generated by the scenario (high mtROS, high HIF-1). (*E*) The fractions of the L/L phenotype corresponding to each scenario in *D*. The analysis was repeated for three times and error bars were added. The color codes for different metabolic states “W”, “O”, “W/O” and “L/L” are consistent in figures *B-D*.

To study how cellular network changes its dynamics under parameter perturbations, we focus on two factors: HIF-1 degradation and mtROS production. These two factors are chosen as cancer cells often face varying hypoxic conditions [22], and also undergo changes in their mitochondria [2]. Previously, we showed that stabilized HIF-1 or elevated production of mtROS promotes the “W/O” state. Here, we focused on their effects on the acquisition of the “L/L” state. To investigate the emergence of the “L/L” state, we considered four scenarios - (high mtROS, high HIF-1), (high mtROS, low HIF-1), (low mtROS, high HIF-1) and (low mtROS, low HIF-1). We utilize a variation of our previously developed computation method called RAndom CIrcuit PErturbation (RACIPE) [23] to identify the robust metabolic states resulting from each scenario. Specifically, we randomly sampled values for each parameter from the range (75%P_0_, 125%P_0_) where P_0_ is the baseline value; this was done for all parameters except for the degradation rate of HIF-1 and production rate of mtROS, which are fixed so as to distinguish among the different scenarios. We construct 500 sets of randomly sampled parameters, calculate the stable steadystate solutions for each set of parameters, and collect solutions from all sets of parameters for statistical analysis by which the dominant metabolic profiles can be identified. We find that the fraction of the “L/L” metabolic state is lowest when both the HIF-1 level and the mtROS level are high (**figure 1D i, electronic supplementary material, figure S2 i**), intermediate when either HIF-1 or mtROS are high (**figures 1D-E ii-iii, electronic supplementary material, figure S2 ii -iii**), and highest when both the HIF-1 level and the mtROS level are low (**figure 1D iv, electronic supplementary material, figure S2 iv**). In summary, our modeling analysis shows that cancer cells can exist in four distinct metabolic states and the occupancies of different metabolic states can be altered by specific kinetic parameter modifications. Specifically, we show that the metabolically L/L state is enriched when the degradation rate of HIF-1 is high or the production rate of mtROS is low.

### Idling Melanoma Cells Exhibit the Metabolic L/L Characteristics

We recently reported that BRAF-mutated SKMEL5 melanoma cells enter a non-quiescent “idling” population state upon long-term MAPKi treatment [19]. Briefly, melanoma cells, including isogenic single-cell derived subclones (SC01, SC07, SC10), exhibit an early differential drug response before transitioning into an idling state characterized by ~zero net-growth [19]. In particular, SC01 and SC10 had divergent initial response to BRAFi (negative for SC01, positive for SC10), but both subclones exhibit ~zero net-growth rates under continued exposure to MAPKi, while SC07 maintained its initial ~zero net-growth (**figure 2A**). At the population level, the idling state exhibits stable population size, realized by an approximately equal rates of division and death [19]. Since recent studies show that melanoma cells treated with MAPKi undergo metabolic reprogramming [4,24], we seek to characterize the metabolic profiles of isogenic subclones at baseline and en route to the idling state.

**Figure 2.**
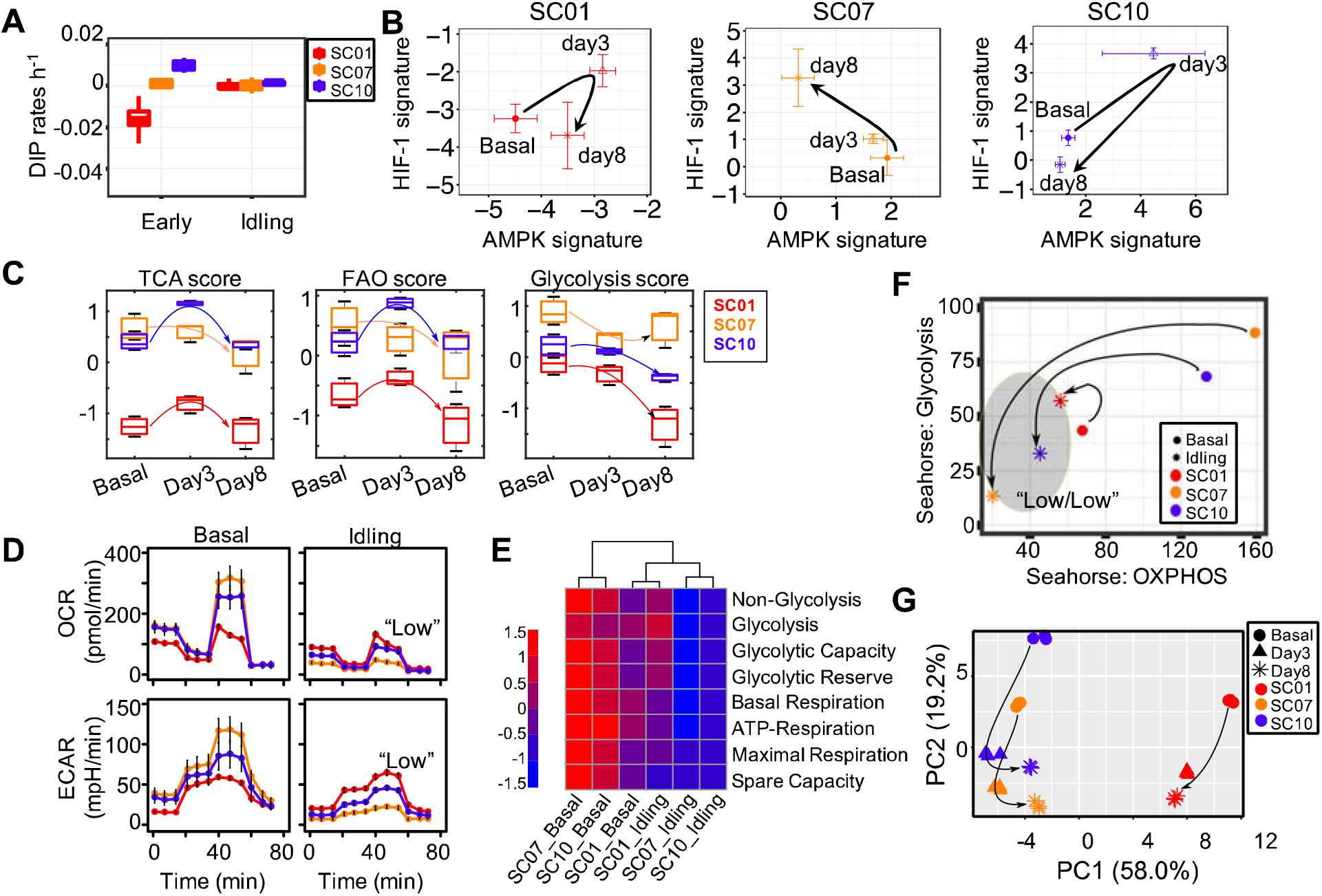
Melanoma cells entering an idling state exhibit the L/L feature. (*A*) Growth rate of three single-cell-derived SKMEL5 subclones, SC01, SC07 and SC10 in either *early* (untreated) condition or in the *Idling* state (8 days in 8uM PLX4720). (*B*) The mean AMPK and HIF-1 signatures of each subclone at indicated time points. (*C*) Box plots for the TCA score, FAO score and glycolysis score of SC01, SC07, and SC10. For SC01: TCA score, P_basal-day3_=0.04; FAO score, P_day3-day8_=0.03; glycolysis score, Pbasal-day8=0.01, P_day3-day8_=0.02. For SC10: TCA score, P_basal-day3_=0.003, P_day3-day8_<0.0001; FAO score, P_basal-day3_=0.009, P_day3-day8_=0.004; glycolysis score, P_basal-day8_=0.01, P_day3-day8_<0.0009. (*D*) OCR (top panel) and ECAR (bottom panel) of three SC01, SC07, and SC10 in either their *basal* (untreated) conditions or in *Idling* state (8 days in 8uM PLX4720), error bars mean ± SEM. The seahorse data is normalized to cell count. (*E*) Heat-map of metabolic parameters for three subclones in *basal* vs *idling* state extracted from Seahorse assays in Figure 2D - the first four parameters from a glycolytic function experiment (Glyco Stress Test) and the second four parameters from a mitochondria function experiment (Mito Stress Test) (**electronic supplementary material, §S2**). (*F*) Aggregated glycolytic parameters plotted against aggregated OXPHOS parameters for three subclones in *basal* vs *idling* state, arrows show the direction of change in idling state from their basal counterparts. (*G*) First two principal components from the time-course RNA-Seq data on genes involved in metabolism in three subclones, SC01, SC07, and SC10 in control, 3 days and 8 days in 8uM PLX4720, arrows indicate the shift in transcriptome of subclones along PC1 and PC2 axes.

To characterize the change of metabolic activity in single-cell derived subclones upon treatment, we apply the previously developed AMPK/HIF-1 signatures and metabolic pathway scores to quantify the activity of AMPK and HIF-1, and the activity of three major metabolic processes - the citric acid cycle (TCA), FAO and glycolysis. Strikingly, by applying these signatures to the longitudinal RNA-Seq data of the isogenic subclones upon BRAFi at baseline (0d), day 3 (3d) and at day 8 (8d, idling state), we find the subclone SC01 exhibit lowest AMPK/HIF-1 activity and lowest TCA/FAO/glycolysis activity (**figures 2B-C**, **electronic supplementary material, figure S3**). Upon MAPKi treatment, we find that the melanoma cells increase both their AMPK and HIF-1 activities from baseline to day 3 and then decrease both from day 3 to day 8, with an overall decrease from day 0 to day 8, especially for the subclones SC01 and SC10 (**figure 2B, electronic supplementary material, figure S3**). The subclone SC07 decreases its AMPK activity while increasing its HIF-1 activity from baseline to day 3 to day 8 (**figure 2B, electronic supplementary material, figure S3**). The TCA/FAO activity for both SC01 and SC10 initially increases and then decreases from day 3 to day 8, with an overall decrease from day 0 to day 8. The glycolysis activity of both SC01 and SC10 decreases from its baseline to day 8 (**figure 2C**). Interestingly, the TCA/FAO activity of SC07 decrease from its baseline to day 8 (**figure 2C**), In the only outlier, the glycolysis score of SC07 first decreases, and then slightly increases. Nonetheless, taken all together our analysis suggests that change in central metabolism has a critical role in drug response.

To further test whether the convergence to the L/L state is a common trend of melanoma cells upon long-term treatment, we apply the AMPK/HIF-1 signatures and the metabolic pathway scores to quantify the change of metabolic profiles of two *BRAF*-mutated melanoma cell lines, M397 and M229 upon vemurafenib treatment for up to three months [25]. Our analyses show that while the M397 and M229 cells have different initial AMPK and HIF-1 activities, both exhibit a significant decrease in their AMPK and/or HIF-1 activities in response to vemurafenib within 21 days (**electronic supplementary material, figure S4A**). Interestingly, the M229 cells increase its AMPK activity after day 21 and recover its AMPK activity comparable to its baseline at day 90 (**electronic supplementary material, figure S4A**). In both cell lines, the TCA and glycolysis scores decrease during the course of the treatment while the FAO score does not show a clear difference from its baseline (**electronic supplementary material, figure S4B**), In summary, the M397 and M229 cells exhibit the L/L gene signatures upon vemurafenib treatment.

To directly test the modeling-predicted and metabolic signatures-characterized L/L state, we measure the metabolic activity of cells in the idling state relative to cells at baseline. We quantify both the oxygen consumption rate (OCR) and extracellular acidification rate (ECAR) (see **Materials and Methods**) in isogenic subclones using the Seahorse extracellular flux analyzer platform (Agilent). At baseline, the OCRs are different in different subclones - SC07 (highest), SC10 (intermediate) and SC01 (lowest). In the idling state, we find that the OCR of both SC07 and SC10 are significantly reduced, while SC01 exhibited little change from its already low initial OCR (**figure 2D, top panel**), which is consistent with the characterization results by the metabolic pathway scores (**figure 2C**). Similar results are obtained for ECAR (**figure 2D, bottom**). From the metabolic parameters extracted from Seahorse assays as previously described [6], we aggregate the glycolysis-related and OXPHOS-related parameters (**figure 2E**), and project subclones at baseline and in idling state into two-dimensional metabolic space. Consistent with earlier results (**figures 2C-D**), SC10 exhibits a significant reduction in both its glycolytic and its OXPHOS profile, while SC01 exhibits relatively little change from its already low baseline (**figure 2E**). For direct comparison between glycolysis and OXPHOS activity in these three subclones, we convert glycolytic and OXPHOS rates to uniform units (JATP: pmol ATP/min) [26]. We found that cells in the idling state exhibit reduced bioenergetic capacity in all three subclones, more pronounced in SC07&10 relative to SC01 (**electronic supplementary material, figure S5**). To examine the temporal evolution to the idling state, we perform principal component analysis (PCA) on the RNA-Seq data of the genes related to metabolism based on the Molecular Signatures Database (MSigDB) for all three subclones (**electronic supplementary material, table S1**). Interestingly, metabolic gene expression profiles of all three subclones converge in both the principal components 1 and 2 (PC1 & PC2) when entering the idling state (**figure 2G**). Taken together, these results suggest that the *BRAF*-mutated melanoma cells, including the isogenic subclones, under MAPKi treatment, tend to decrease both their glycolysis and OXPHOS activities and acquire a L/L state, while converging to an idling state. The L/L state thus appears to be a common feature of idling melanoma cells, that could also explain a recently described treatment-induced dedifferentiation in melanoma cells [25].

### Residual Melanoma Tumors on MAPKi Therapy Display the Metabolically L/L Signature

The modeling-predicted and experiment-validated metabolically-inactive L/L state of idling melanoma cells motivates us to further analyze whether such a L/L phenotype can be observed in melanoma patient samples. We analyze the change of the metabolic activity in a cohort of melanoma patients before and during MAPKi treatment [27]. Strikingly, even though there are only 7 patients, we observe a consistent temporal decrease of both AMPK and HIF-1 activity upon MAPKi treatment in 6 of the 7 patients (**figure 3**). Additionally, using the metabolic pathway scores, we observe that the samples that underwent longer treatment (patients 1, 3 and 4) exhibit a decrease of both glycolysis and TCA scores (Figure 3 left panel), while the samples with shorter treatment (patients 6, 7 and 8) exhibit a decrease in glycolysis score (Figure 3 right panel). Patient 2, however, shows a decrease in HIF-1 activity and an increase in AMPK activity (**electronic supplementary material, figure S6A**). In contrast, the FAO score increases upon treatment in almost all patients (**electronic supplementary material, figure S6B**), which suggests that the drug-tolerant cells tend to depend on lipid metabolism as reported before [25]. Consistent with the cell line data, the HIF-1 and AMPK activities of melanoma samples are different at baseline among patients. Interestingly, in almost all cases, there is a decrease in their activities upon treatment (**electronic supplementary material, figure S6C**), indicating a significant trend towards the L/L state.

**Figure 3.**
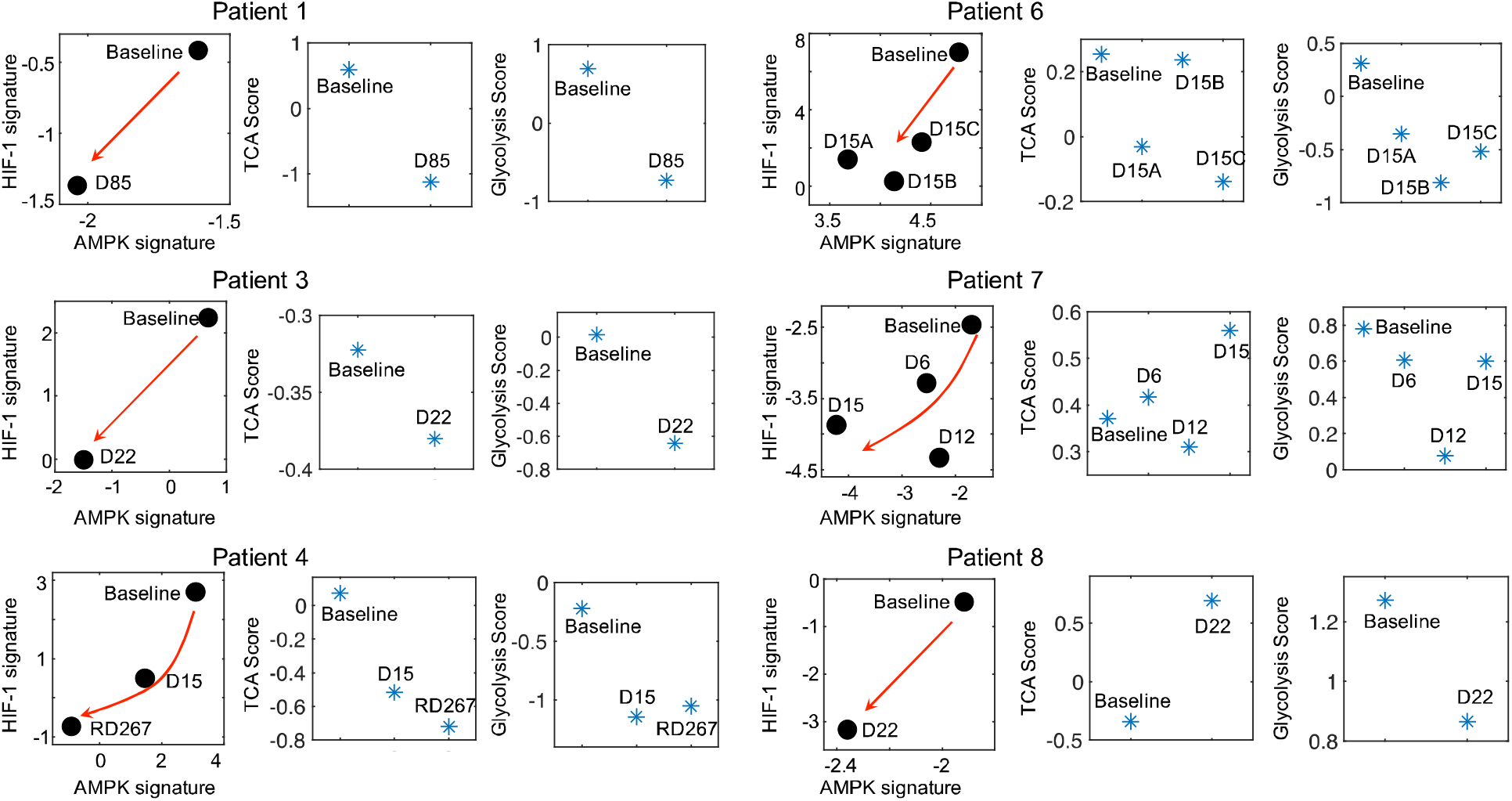
Residual melanoma tumors upon MAPKi therapy exhibit L/L signature. The AMPK/HIF-1 signatures and TCA/glycolysis scores of BRAF-mutated tumor samples upon MAPKi treatment for the indicated time points. The RNA-Seq data are obtained from GEO with the series ID GSE75299. The red arrows denote the direction of the change of AMPK and HIF-1 activity of samples upon MAPKi treatment.

We also obtain the RNA-Seq data of melanoma patient samples from The Cancer Genome Atlas (TCGA) and evaluate their AMPK/HIF-1 activity. We find that the melanoma patient samples can be grouped into four clusters based on their HIF-1 and AMPK signatures (**electronic supplementary material, figure S7**). In addition to patient samples that exhibit either high HIF-1 activity (corresponding to a glycolysis state) or high AMPK activity (corresponding to an OXPHOS state), high activity of both HIF-1 and AMPK (corresponding to a hybrid metabolic state), there exists a fraction of samples that shows low activity of both HIF-1 and AMPK, tentatively characterized as the L/L state (**electronic supplementary material, figure S7**). In summary, our analyses suggest that L/L phenotype is clinically relevant in *BRAF*-mutated melanomas.

## Discussion

Drug-tolerance remains a challenge in anti-cancer therapies. In recent years, metabolic plasticity has been linked to drug-tolerance. However, it remains largely unknown how cancer cells adjust their metabolic profiles to evade treatment. Through coupling gene regulation with metabolic pathways, our metabolic modeling framework characterizes distinct metabolic states in cancer cells [3]. In particular, our model predicts the existence of a metabolically inactive phenotype L/L characterized by low AMPK/HIF-1 activity and low OXPHOS/glycolysis pathway activity. Although metabolic adaptation has been reported in different cancer types [2,4,5,8,28–30], such a L/L metabolic phenotype has not been previously described.

The recently reported idling state acquired by melanoma cells upon long-term MAPKi treatment [19] represents a novel drug-tolerant mechanism that interferes with successful treatment. Notably, the idling state differs from the slow-cycling phenotype [14], and the neural-crest phenotype [18] as described in melanoma resistance. In the present study, we show that these idling melanoma cells exhibit the theory-predicted L/L phenotype characterized by low AMPK/HIF-1 activity and low OXPHOS/glycolysis pathway activity. We further show that melanoma cells tend to decrease their AMPK/HIF-1 activity and TCA/glycolysis activity upon MAPKi treatment and converge toward a L/L phenotype, through analyzing the RNA-Seq data of both cell lines and patient samples obtained from GEO. Taken together, our results suggest that the acquisition of a L/L state tends to be a general trend for melanoma cells upon long-term MAPKi treatment. Future work should focus on elucidating the mechanisms by which melanoma cells acquire the L/L phenotype and how that gives rise to drug-tolerance.

It is worth noting that the acquisition of a L/L phenotype by melanoma cells upon long-term MAPKi treatment is not always direct. Our results suggest that the paths to a L/L phenotype can vary. While the subclones SC01 and SC10 show an initial increase of both their AMPK/HIF-1 and TCA/FAO activities before converging to the L/L phenotype upon the MAPKi treatment, the subclone SC07 continuously decreases its AMPK and TCA/FAO activities. Similarly, in the clinical context, not all patient samples continuously decrease their glycolysis and OXPHOS activity upon treatment. Six out of seven patients did show a decrease in both HIF-1 and AMPK activity; one remaining sample (Patient 2) shows a decrease only in its HIF-1 activity. Notably, the melanoma samples from patients 1, 3 and 4 exhibit more pronounced decrease in both their AMPK/HIF-1 activity and TCA/glycolysis activity upon treatment relative to the samples from patients 6, 7 and 8. One possible explanation is that the samples from patients 1, 3 and 4 were obtained upon longer treatment (85 days, 22 days and 267 days respectively) relative to the samples from patients 6, 7 and 8 (15 days, 15 days and 22 days respectively) (**Figure 3**), as the acquisition of a L/L state may require a long-term treatment. In addition, the differences in treatment regimens may play a role in the acquisition of the L/L phenotype as well as interpatient and inter-tumor heterogeneity. Interestingly, the FAO scores of samples from patients 3, 4, 7, 8 increase upon treatment, which is consistent with the report that these cells may exhibit increased dependence on lipid peroxidation pathways, and hence vulnerable to ferroptosis-inducing drugs [25]. Therefore, it is critical to consider the dynamics of cellular response. It may be necessary to evaluate the change of both genes and the metabolic fluxes to fully characterize the change of metabolic activity in melanoma cells. Indeed, our experimental result (**electronic supplementary material, figure S8**) and recent reports [4,5,13,31] suggest that an increase in OXPHOS under perturbations in cancer cells might rather be an intermediate response, but not a final stable state, as melanoma cells have been shown to undergo drug-induced dedifferentiation [25]. That being said, MAPKi treatment might not always induce a L/L phenotype due to preexisting heterogeneity of melanoma cells. Since tumor metabolism can dynamically change under drug exposure, we cannot rule out the possibility of co-existence of multiple metabolic phenotypes in melanoma cells, the proportion of which depends on cell types, drug treatment, and the time that has elapsed since drug exposure.

As the L/L phenotype potentially allows for tumor relapse, it is critical to identify the conditions that promote its existence and enrichment. Using our metabolic modeling framework, we show that high degradation rate of HIF-1 and/or low production rate of mtROS can significantly enhance the L/L phenotype. This is consistent with recent reports that show BRAFi treatment induces degradation of HIF-1α at both mRNA and protein levels in drug-tolerant *BRAF*-mutated melanoma cells [7,30]. Additional experimental work to characterize the change of HIF-1 and mtROS in idling cells relative to non-idling cells needs to be done to test this prediction.

The L/L phenotype is potentially observed in a fraction of patient samples that exhibit low activity of both AMPK and HIF-1. Since treatment information regarding patient samples in TCGA is incomplete, it is hard to evaluate the effect of treatment on the acquisition of the L/L phenotype. We speculate that the L/L phenotype enables cancer cells to maintain cell viability and, because of the ongoing cell division, to accumulate mutations *en route* to resistance. One could also speculate that repression of central-carbon metabolism (i.e., decreased glycolysis and OXPHOS activity), as we see in the idling cells, allows cells to synthesize biomolecules to maintain cellular homeostasis [11]. This is consistent with recent reports which suggest that drug-tolerant cancer cells exhibit an increased dependence on polyunsaturated lipids for survival [8]. The AMPK/HIF-1 signatures and metabolic pathway scores can not only capture the trajectory of treatment-induced metabolic changes but also delineate the differences among different cells and different patient samples. Therefore, the AMPK/HIF-1 signatures may be used to further explore the existence of a L/L phenotype in other types of cancer, such as breast cancer [3]. This will eventually lead to the understanding of how L/L phenotype could be exploited therapeutically.

There are several limitations about the present study. First, without including the details of the MAPK/ERK pathway, our current mathematical model cannot mechanistically explain the heterogeneous initial responses of SC01, 07 and 10 to MAPKi treatment. Second, to explore whether the convergence to a L/L phenotype is a general feature of melanoma cells, both metabolic activity and fluxes upon MAPKi treatment need to be performed for multiple melanoma cells. Third, our metabolism model does not explicitly focus on the glutamine pathway. We are assuming an available supply of this needed input and that this supply is used for biosynthesis of e.g. fatty acids that are needed for cell growth which as already mentioned is non-negligible in the idling state. Future work that aims to understand in more detail the difference in division rates between cells in the idling population and slow-cycling cells on the one hand versus untreated melanoma cells on the other hand will involve an extension of our model to include glutamine intake and utilization.

The heterogeneity in melanomas might result in differential proportions of various metabolic phenotypes that can co-exist. Initial proportions of cancer cells in these metabolic states may affect how they respond to perturbations, as indicated by a three-state (OXPHOS^High^Glycolysis^High^ (H), OXPHOS^Low^Glycolysis^Low^ (L), OXPHOS^High^Glycolysis^Low^ or OXPHOS^Low^Glycolysis^High^ (Intermediate, I)) cell population model (**electronic supplementary material, §S3 and figures S9-10**). For simplicity, we grouped OXPHOS^High^Glycolysis^Low^ and OXPHOS^Low^Glycolysis^High^ as intermediate metabolic state. The model assumes that the growth rate is a function of cellular capacity to utilize OXPHOS and glycolysis. This three-state model can qualitatively reproduce the different drug responses of SC01, 07 and 10 and also sheds light on the paths towards convergence to low-low metabolic state. This path becomes especially useful when the perturbations affect one or the other metabolic pathways. For instance, if cancer cells undergo metabolic reprogramming via a decrease in their mitochondrial respiration, they may not respond to OXPHOS inhibitors. Similarly, if cancer cells re-route their metabolic needs via a decrease in their glycolytic capacity, they may not respond to glycolytic inhibitors. The resulting intermediate state that cells acquire during reprogramming might expose vulnerabilities that can be therapeutically targeted [32] (**figure 4**). Therefore, metabolic heterogeneity should be quantified as distinct phenotypes to examine the initial tumor composition, and their functional significance, and studied dynamically to decipher the likely escape route for a rational combination therapy (**figure 4**). Detailed mapping of the cellular bioenergetics to quantify the contribution of the energy derived from metabolic processes to either cell division or cell death is outside the scope of this paper, and is an active area for future investigation.

**Figure 4.**
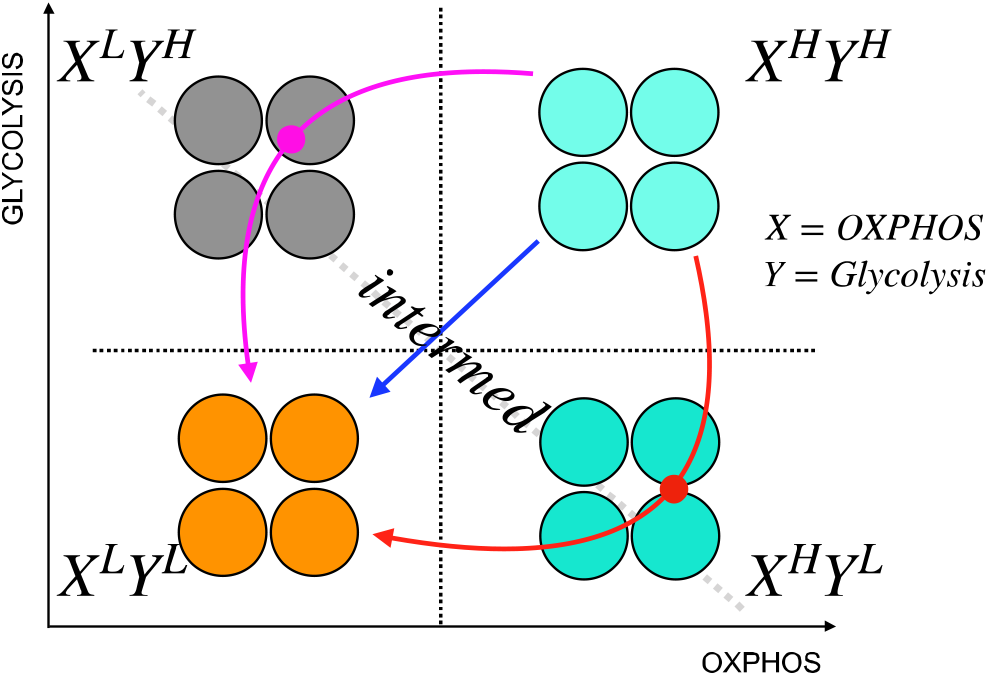
A schematic illustration of cancer metabolic plasticity and resulting phenotypes. Tumor populations can exhibit distinct metabolic phenotypes that co-exist, and the proportions of cells in each state give rise to unique cell phenotype, and metabolic profiles. Depending on the cell types, and the drugs used, tumor cells could follow different paths to switch among the phenotypic states. In addition, tumor cells can transition into the same end state (L/L) via different paths; the red path shows the transition of tumor cells from X^High^Y^High^ state to X^Low^Y^Low^ state via X^high^Y^low^ state, the magenta path shows the transition from X^High^Y^High^ state to X^Low^Y^Low^ state via X^Low^Y^high^ state, and the blue path shows a direct transition from X^High^Y^High^ state to X^Low^Y^Low^ state. In this schematic, X represents capacity of cells to use OXPHOS, and Y represents the capacity of cells to use glycolysis.

Abnormal metabolism has been observed in cancer for over a century. For quite some time, aerobic glycolysis was regarded as the dominant metabolic phenotype. It was hypothesized that the elevated glycolytic activity in cancer cells was due to the defects in their mitochondria. Recently, it has been recognized that cancer mitochondria are not, in general, dysfunctional but instead they are actively functioning in certain cancer types. By performing systems biology analysis of cancer metabolism, we show that cells are able to acquire a hybrid metabolic phenotype, and a metabolically L/L phenotype in addition to glycolysis and OXPHOS. From the Warburg effect, to OXPHOS, to hybrid, and now to the L/L phenotype, we are getting closer to a complete picture of the possible metabolic states acquired by cancer cells and how cancer cells adjust their metabolic profiles under perturbations. Since recent efforts in targeting cancer metabolism have been largely ineffective, a better understanding of cancer metabolic plasticity would eventually contribute to more effective therapeutic strategies. We foresee the integration of computational and experimental approaches will be necessary to elucidate mechanisms underlying cancer cell behaviors. Therefore, future studies aimed at limiting metabolic plasticity, metabolic reverse engineering, may provide insights into novel therapeutic regimens.

## Acknowledgments

We would like to thank Prof. Kevin Janes (University of Virginia) and Prof. Benny Abraham Kaipparettu (Baylor College of Medicine, Houston, TX) for critical reviews of the manuscript.

## References

1. Vander Heiden MG, Cantley LC, Thompson CB. 2009 Understanding the Warburg effect: the metabolic requirements of cell proliferation. Science 324, 1029–1033.

2. Porporato PE et al. 2014 A mitochondrial switch promotes tumor metastasis. Cell Rep. 8, 754–766.

3. Jia D, Lu M, Jung KH, Park JH, Yu L, Onuchic JN, Kaipparettu BA, Levine H. 2019 Elucidating cancer metabolic plasticity by coupling gene regulation with metabolic pathways. Proc. Natl. Acad. Sci. U. S. A. 116, 3909–3918.

4. Haq R et al. 2013 Oncogenic BRAF regulates oxidative metabolism via PGC1α and MITF. Cancer Cell 23, 302–315.

5. Baenke F et al. 2016 Resistance to BRAF inhibitors induces glutamine dependency in melanoma cells. Mol. Oncol. 10, 73–84.

6. Hardeman KN, Peng C, Paudel BB, Meyer CT, Luong T, Tyson DR, Young JD, Quaranta V, Fessel JP. 2017 Dependence On Glycolysis Sensitizes BRAF-mutated Melanomas For Increased Response To Targeted BRAF Inhibition. Sci. Rep. 7, 42604.

7. Parmenter TJ et al. 2014 Response of BRAF-mutant melanoma to BRAF inhibition is mediated by a network of transcriptional regulators of glycolysis. Cancer Discov. 4, 423–433.

8. Viswanathan VS et al. 2017 Dependency of a therapy-resistant state of cancer cells on a lipid peroxidase pathway. Nature 547, 453–457.

9. Wood TK, Knabel SJ, Kwan BW. 2013 Bacterial Persister Cell Formation and Dormancy. Applied and Environmental Microbiology. 79, 7116–7121. (doi:10.1128/aem.02636-13)

10. Sharma SV et al. 2010 A chromatin-mediated reversible drug-tolerant state in cancer cell subpopulations. Cell 141, 69–80.

11. Collado M, Blasco MA, Serrano M. 2007 Cellular senescence in cancer and aging. Cell 130, 223–233.

12. Aguirre-Ghiso JA. 2007 Models, mechanisms and clinical evidence for cancer dormancy. Nat. Rev. Cancer 7, 834–846.

13. Ravindran Menon D et al. 2015 A stress-induced early innate response causes multidrug tolerance in melanoma. Oncogene 34, 4448–4459.

14. Ahn A, Chatterjee A, Eccles MR. 2017 The Slow Cycling Phenotype: A Growing Problem for Treatment Resistance in Melanoma. Mol. Cancer Ther. 16, 1002–1009.

15. Perego M et al. 2018 A slow-cycling subpopulation of melanoma cells with highly invasive properties. Oncogene 37, 302–312.

16. Müller J et al. 2014 Low MITF/AXL ratio predicts early resistance to multiple targeted drugs in melanoma. Nat. Commun. 5, 5712.

17. Konieczkowski DJ et al. 2014 A melanoma cell state distinction influences sensitivity to MAPK pathway inhibitors. Cancer Discov. 4, 816–827.

18. Su Y et al. 2017 Single-cell analysis resolves the cell state transition and signaling dynamics associated with melanoma drug-induced resistance. Proc. Natl. Acad. Sci. U. S. A. 114, 13679–13684.

19. Paudel BB, Harris LA, Hardeman KN, Abugable AA, Hayford CE, Tyson DR, Quaranta V. 2018 A Nonquiescent ‘Idling’ Population State in Drug-Treated, BRAF-Mutated Melanoma. Biophys. J. 114, 1499–1511.

20. Yu L, Lu M, Jia D, Ma J, Ben-Jacob E, Levine H, Kaipparettu BA, Onuchic JN. 2017 Modeling the Genetic Regulation of Cancer Metabolism: Interplay between Glycolysis and Oxidative Phosphorylation. Cancer Res. 77, 1564–1574.

21. Meyer CT et al. 2019 Quantifying Drug Combination Synergy along Potency and Efficacy Axes. Cell Syst 8, 97–108.e16.

22. Eales KL, Hollinshead KER, Tennant DA. 2016 Hypoxia and metabolic adaptation of cancer cells. Oncogenesis 5, e190.

23. Huang B, Lu M, Jia D, Ben-Jacob E, Levine H, Onuchic JN. 2017 Interrogating the topological robustness of gene regulatory circuits by randomization. PLoS Comput. Biol. 13, e1005456.

24. Abildgaard C, Guldberg P. 2015 Molecular drivers of cellular metabolic reprogramming in melanoma. Trends Mol. Med. 21, 164–171.

25. Tsoi J et al. 2018 Multi-stage Differentiation Defines Melanoma Subtypes with Differential Vulnerability to Drug-Induced Iron-Dependent Oxidative Stress. Cancer Cell 33, 890–904.e5.

26. Mookerjee SA, Gerencser AA, Nicholls DG, Brand MD. 2018 Quantifying intracellular rates of glycolytic and oxidative ATP production and consumption using extracellular flux measurements. J. Biol. Chem. 293, 12649–12652.

27. Song C, Piva M, Sun L, Hong A, Moriceau G, Kong X. 2017 Recurrent tumor cell–intrinsic and–extrinsic alterations during MAPKi-induced melanoma regression and early adaptation. Cancer Discov.

28. Hangauer MJ et al. 2017 Drug-tolerant persister cancer cells are vulnerable to GPX4 inhibition. Nature 551, 247–250.

29. Vazquez F et al. 2013 PGC1α expression defines a subset of human melanoma tumors with increased mitochondrial capacity and resistance to oxidative stress. Cancer Cell 23, 287–301.

30. Cesi G, Walbrecq G, Zimmer A, Kreis S, Haan C. 2017 ROS production induced by BRAF inhibitor treatment rewires metabolic processes affecting cell growth of melanoma cells. Mol. Cancer 16, 102.

31. Smith MP et al. 2016 Inhibiting Drivers of Non-mutational Drug Tolerance Is a Salvage Strategy for Targeted Melanoma Therapy. Cancer Cell 29, 270–284.

32. Paudel BB, Quaranta V. 2019 Dynamics of drug response informs rational combination regimens. Sci. Signal. 12. (doi:10.1126/scisignal.aax9742)

